# The Meatball Matchup: Plant vs. Animal Proteins on Campus

**DOI:** 10.64898/2026.03.05.709981

**Authors:** Skyler R. St. Pierre, Aeneas O Koosis, Nancy Zhang, Ellen Kuhl

## Abstract

Institutional dining halls increasingly serve plant-based meat products, yet it remains unclear how consumers perceive these products relative to animal-based options in real-world settings. This study compares consumer sensory perceptions of two plant-based meatballs (soy, soy-wheat) and two animal-based meatballs (beef, beef–mushroom) among university dining hall patrons (*n* = 116), complemented by instrumental Texture Profile Analysis. Animal-based meatballs received significantly higher ratings for moistness, meatiness, fattiness, and tastiness (all *p* < 0.001), with the meatiness gap being the largest (Δ = 1.40 on a 5-point scale). Texture analysis revealed that animal-based samples were significantly harder, more cohesive, and chewier than plant-based samples. In contrast, consumers perceived no differences in chewiness or hardness between categories, revealing a disconnect between instrumental and sensory measures. Just-About-Right penalty analysis identified insufficient savoriness as a universal improvement target across all products, including beef. Flavor and texture emerged as the dominant drivers of dining choice, while sustainability and animal welfare ranked lowest in importance. These findings suggest that sensory parity—particularly in moistness, meatiness, and savoriness—may drive acceptance of plant-based meat more strongly than sustainability messaging. Data and code are available at https://github.com/LivingMatterLab/AI4Food

## 1. Introduction

The global plant-based meat market was valued at approximately 6.1 billion (USD) in retail sales in 2024, with the United States representing the largest single market at $1.2B (Good Food Institute, 2025). Despite this substantial market size, plant-based meat and seafood account for only 1.7% of total US retail packaged meat dollar sales, and household penetration has stabilized at approximately 13% (Good Food Institute, 2025). US retail sales declined 7% in 2024 (SPINS Retail Data, 2025), raising questions about whether sensory quality limitations are constraining broader adoption (Peacock, 2026).

Plant-based meat alternatives are designed to replicate the sensory experience of conventional meat products, yet achieving sensory parity remains a significant technical challenge (He et al., 2020; Kumari et al., 2024). Plant-based products are frequently perceived as failing to meet the expectations set by “tastes like meat” marketing, particularly among flexitarian consumers who still consume traditional animal-based foods (Fiorentini et al., 2023). The sensory gap is especially pronounced for texture (van den Bedem et al., 2026): plant-based products are generally softer and less chewy than animal-derived meats, and replicating the fibrous, anisotropic protein structure of muscle tissue remains difficult using plant proteins (McClements & Grossmann, 2021; St. Pierre & Kuhl, 2024a).

University dining halls represent a strategically important context for plant-based meat adoption. Approximately 14% of Gen Z consumers actively limit meat consumption, and young adults aged 18–24 demonstrate higher openness to plant-based alternatives compared to older demographics (Bryant, 2019; Green et al., 2026). However, most existing sensory research on plant-based meat has been conducted in controlled laboratory conditions rather than real-world dining environments (Fiorentini et al., 2023; Bakhsh et al., 2021). Additionally, all-you-can-eat meal plans remove fiscal barriers to choosing healthier, more expensive food options (Strong et al., 2008).

Texture is a critical determinant of meat acceptability (Kuhl, 2025). Texture Profile Analysis (TPA) provides objective instrumental measures of mechanical properties including hardness, cohesiveness, springiness, and chewiness (Bourne, 2002). Plant-based meat alternatives consistently exhibit lower hardness and chewiness than animal-derived counterparts in TPA studies, but it is unclear to what extent these instrumental differences translate to consumer perception (Bakhsh et al., 2021; Kamani et al., 2023). Using small untrained tasting groups (*n* = 16-18), we have previously found that sensory brittleness is correlated with mechanical stiffness in deli slices (St. Pierre et al., 2025) and sensory hardness is associated with mechanical stiffness in minced meat products (St. Pierre et al., 2024b) and fungi-based steaks (Vervenne et al., 2025). Understanding the relationship between instrumental texture properties and consumer perceptions is essential for guiding product development (Datta et al., 2026).

The Just-About-Right (JAR) scale methodology identifies specific product attributes requiring reformulation by quantifying the impact of attribute deviations on overall liking (Rothman & Parker, 2009). This diagnostic approach is particularly valuable for comparing multiple products and prioritizing improvement opportunities based on actionable penalties. Consumer food choices are influenced by multiple factors beyond sensory properties, including health perceptions, environmental concerns, familiarity, and social context (Onwezen et al., 2021). Students in university dining settings respond more favorably to plant-forward dishes presented by taste-focused labels rather than health-focused or basic labels (Turnwald et al., 2019). The relative importance of hedonic versus ethical factors in real-world dining hall settings has not been systematically quantified.

*The objective of this study is to quantify sensory and mechanical gaps between plant-based and animal-based meatballs and identify attributes that limit consumer adoption*.

We compared consumer sensory perceptions of plant-based and animal-based meatballs in a university dining setting, characterized instrumental texture properties using TPA, and examined their correspondence with consumer perception. We then identified product-specific reformulation priorities through JAR penalty analysis and assessed the relative importance of hedonic, health, and ethical factors in dining hall food choices.

## 2. Materials and Methods

### 2.1. Participants

We recruited participants from a dining hall at Stanford University during regular meal service. Inclusion criteria were: i) age 18 years or older, ii) current dining hall access, and iii) no allergies to soy or pea protein. Stanford University Institutional Review Board approved the study (eProtocol #82951) classified as exempt under taste and food quality evaluation (45 CFR 46.104(d)(6)). Participants provided electronic informed consent via Qualtrics and received a $10 gift card upon survey completion.

### 2.2. Meatball Products

We evaluated four meatball products, two plant-based formulations labeled as *Soy* and *Soy-Wheat*, and two animal-based formulations labeled as *Beef* and *Beef-Mushroom* (Table 1). The Soy and Soy-Wheat meatballs represent leading commercial brands with different protein bases. The Beef-Mushroom meatball is prepared in-house by the Stanford Dining Halls and commonly served on campus.

**Table 1:**
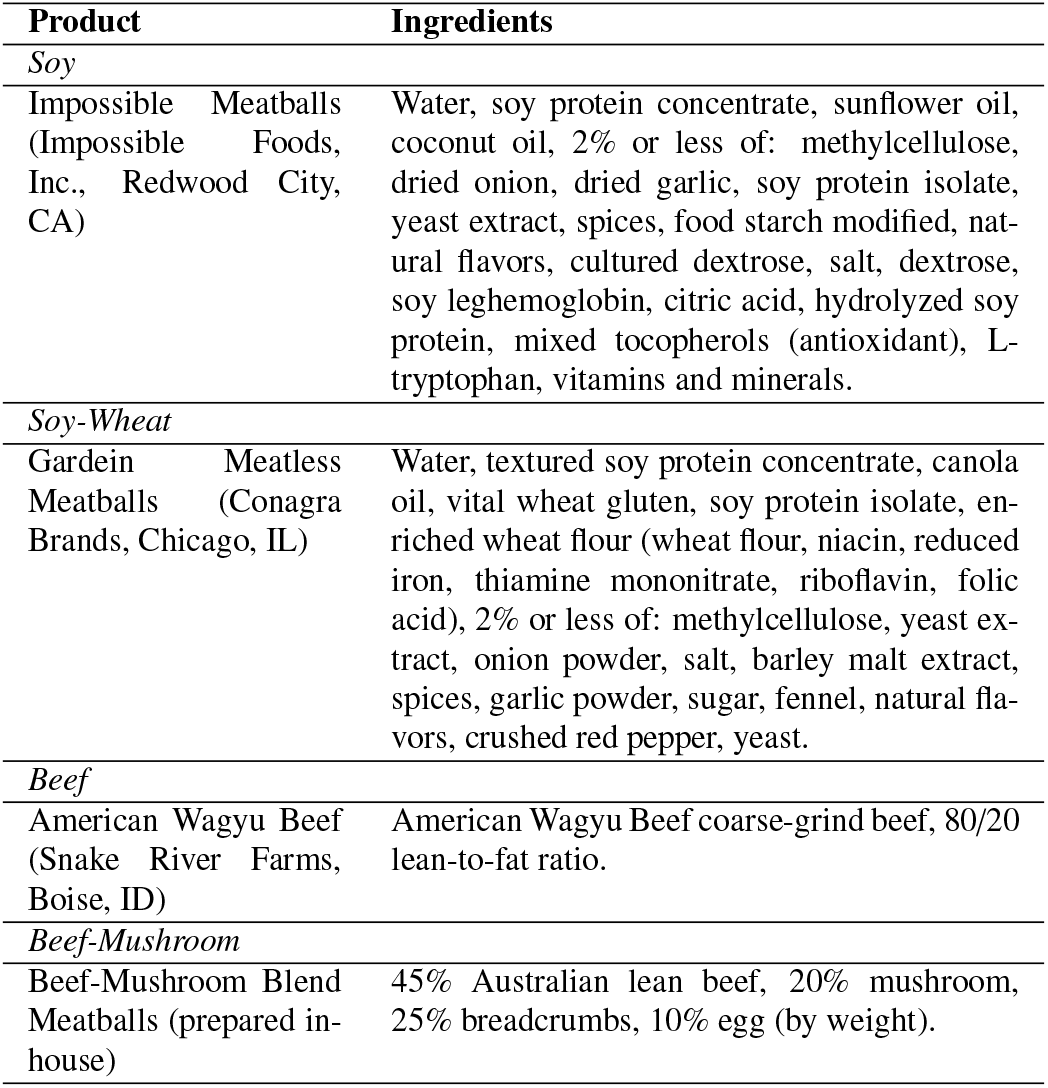
Meatball products. This study evaluated two plant-based and two animal meatballs listed with product information and ingredients.

### 2.3. Sample Preparation

Culinary staff prepared all meatballs according to manufacturer package instructions (plant-based products) or standard institutional preparation protocols (beef products). Plant-based meatballs were baked at 190^°^C for 18–20 minutes. Beef meatballs were formed to approximately 30 g portions and baked at 180^°^C until internal temperature reached 74^°^C. Samples were served warm and offered with optional pasta accompaniments, consistent with typical dining hall presentation (Figure 1, left).

**Figure 1:**
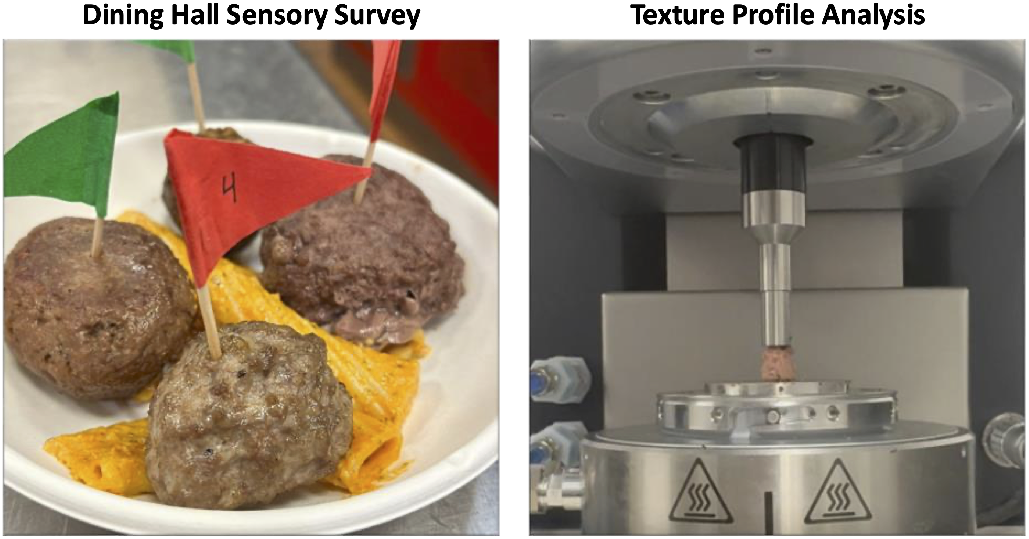
Dining hall sensory survey and texture profile analysis. Left: Four meatball products with a small serving of pasta and labeled with flags for Soy (1), Soy-Wheat (2), Beef (3), and Beef-Mushroom (4) meatballs. Right: A cylindrical 8 mm by 10 mm meatball sample during double-compression testing in the rheometer.

We labeled all meatballs with colored and numbered flags to comply with standard university labeling of vegetarian/vegan dishes and to facilitate rapid product identification while maintaining brand anonymity. We also displayed all allergens.

### 2.4. Consumer Sensory Evaluation

#### 2.4.1. Procedure

We collected data during a single lunch service period. We kept all four meatball samples on warming plates and served them simultaneously in a single bowl with a small amount of pasta in addition to regular dining options. We invited participants who selected the meatball samples to participate via table signage that displayed a QR code linked to the survey. The survey proceeded in the following order: (1) demographics and dietary habits, (2) Meat Attachment Questionnaire, (3) sequential sensory evaluation of each meatball with attribute ratings and JAR scales, (4) dining hall choice factor importance ratings, and (5) open-ended feedback. Participants completed the survey on personal mobile devices while consuming the samples. A total of *n* = 116 participants completed the survey for all four meatballs.

#### 2.4.2. Sensory Attribute Ratings

Participants rated each meatball on eight sensory attributes using 5-point category scales (Lawless & Heymann, 2010): chewiness (1 = breaks down instantly; 5 = requires repeated chewing), hardness (1 = extremely soft; 5 = very hard), moistness (1 = very dry; 5 = very juicy), fibrousness (1 = completely uniform; 5 = clear directional fibers), meatiness (1 = no resemblance to meat; 5 = very close to real meat), fattiness (1 = not fatty at all; 5 = very fatty), tastiness (1 = very unpleasant; 5 = delicious), and softness (1 = very hard; 5 = extremely soft). We selected these attributes in line with prior literature that identified key textural attributes (Szczesniak, 2002; Nishinari & Fang, 2018), key sensory attributes (Onwezen et al., 2021), and overall hedonic response such as tastiness.

#### 2.4.3. Just-About-Right Analysis

For each meatball, participants rated five attributes on 3-point Just-About-Right scales: moistness, chewiness, savoriness, fattiness, and fibrousness (1 = not enough, 2 = just about right, 3 = too much). We selected a 3-point scale, because it directly identifies actionable product improvement directions (too much/not enough), and its simplicity reduces respondent burden in a real-world dining context.

#### 2.4.4. Meat Attachment Questionnaire

We assessed psychological attachment to meat using the 5-item subset of the Meat Attachment Questionnaire (MAQ) focused on dependence on meat (Graça et al., 2015). The questions are: 1) I don’t picture myself without eating meat regularly, 2) if I couldn’t eat meat I would feel weak, 3) I would feel fine with a meatless diet [reverse-scored], 4) if I was forced to stop eating meat I would feel weak, and 5) meat is irreplaceable in my diet. Items were rated on 5-point Likert scales (1 = strongly disagree; 5 = strongly agree). The composite score was computed as the mean of all five items.

#### 2.4.5. Choice Factor Importance

Participants rated the importance of eight factors in their typical dining hall food choices using visual analog sliders (0–100): flavor, texture, health/nutrition, social/cultural norms, climate sustainability, animal welfare, familiarity/comfort, and availability. We adapted these factors from prior research on plant-based meat choice motivation and tailored them to a university dining hall context (Estell et al., 2021; Onwezen et al., 2021; Szenderék et al., 2022).

### 2.5. Texture Profile Analysis

We prepared *n* = 10 samples per meatball type. We extracted the cores of each meatball and used an 8mm diameter biopsy punch to create a total of *n* = 40 cylindrical samples with approximately 10 mm height. Instrumental texture analysis was conducted using a Discovery HR 20 Rheometer (TA Instruments, New Castle, DE) (Figure 1, right). Cooked meatballs were equilibrated to room temperature (22 ± 1^°^C) prior to analysis. Each sample was compressed twice to 50% of its original height at a strain rate, 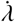, of 25%/s in loading and unloading resulting in a total test time of 8 s.

From the recorded force-time curves, we calculated six TPA parameters: stiffness [N/mm], hardness [N], cohesiveness, springiness, resilience, and chewiness [N] (St. Pierre & Kuhl, 2024a). Three samples were excluded from analysis due to slip-page during testing. Following standard definitions (Bourne, 1978; Friedman et al., 1963), we denote the peak forces of the first and second loading cycles as *F*_1_ and *F*_2_, the associated loading times as *t*_1_ and *t*_2_, the areas under their loading paths as *A*_1_ and *A*_3_, and the areas under their unloading paths as *A*_2_ and *A*_4_ (Dunne et al., 2025). From these characteristic values, we extract six texture profile analysis parameters (Nishinari et al., 2013): The stiffness *E* = σ/ε refers to the slope of the stress-strain curve during the first compression. The hardness *F*_1_ is the peak force during the first compression cycle. The cohesiveness (*A*_3_ + *A*_4_)/(*A*_1_ + *A*_2_) characterizes the material integrity during the second loading and unloading cycle compared to the first. The springiness *t*_2_/*t*_1_ describes how much the material springs back to its original state after the second cycle compared to the first. The resilience *A*_2_/*A*_1_ measures how well a sample recovers between the first unloading and loading paths. The chewiness *F*_1_ (*A*_3_ + *A*_4_)/(*A*_1_ + *A*_2_) *t*_2_/*t*_1_, is the product of hardness, cohesiveness, and springiness.

### 2.6. Statistical Analysis

We conducted all analyses using custom Python scripts. We set the significance threshold to α = 0.05 for omnibus tests. For post-hoc pairwise comparisons, we applied Bonferroni correction (adjusted α = 0.0083 for 6 comparisons). We used Friedman’s test for repeated-measures comparisons across the four meatballs and reported the effect size as Kendall’s *W*. Pairwise comparisons used Wilcoxon signed-rank tests with effect size 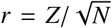. Between-group comparisons used Mann-Whitney *U* tests (2 groups) or Kruskal-Wallis *H* tests (>2 groups). We used Kendall’s *τ* for correlations. Mean-drop analysis followed industry-standard protocols (Rothman & Parker, 2009), with penalties ≥0.50 on the 5-point Tastiness scale flagged as actionable. We used one-way ANOVA with Tukey HSD post-hoc tests to compare TPA parameters across the four products.

We used penalty analysis to quantify the impact of attribute deviations on overall liking by calculating the mean drop in the dependent variable, tastiness, when an attribute deviates from “just about right”. For each attribute, the penalty equals the difference between the mean tastiness rating among respondents who rated the attribute “just about right” and the mean tastiness rating among those who rated it “not enough” or “too much.”

Larger penalties indicate that the deviation more strongly diminishes overall product acceptance, and make that attribute a higher priority for reformulation. We defined actionable deviations as penalties ≥0.50 on the 5-point tastiness scale with *p* < 0.05, following standard thresholds (Rothman & Parker, 2009).

## 3. Results

### 3.1. Participant Characteristics

Table 2 reveals demographic information and dietary habits of the participants in our dining hall study. The sample was pre-dominantly young adults aged 18-24 (96%) with balanced gender representation. Most participants identified as omnivores (94%). Notably, our participants had a high average Meat Dependence score, 4.03/5, indicating a strong dependence on meat in their diets.

**Table 2:** Participant demographics. The study population were predominantly young adults with balanced gender representation (*n* = 116).

| Characteristic | n | % | Characteristic | n | % |
| --- | --- | --- | --- | --- | --- |
| *Age* |  |  | *Gender* |  |  |
| 18–24 | 111 | 95.7 | Male | 60 | 51.7 |
| 25–34 | 1 | 0.9 | Female | 55 | 47.4 |
| > 34 | 4 | 3.4 | Non-binary | 1 | 0.9 |
| *Ethnicity* |  |  | *Diet Type* |  |  |
| Asian | 53 | 45.7 | Omnivore | 109 | 94.0 |
| White | 20 | 17.2 | Flexitarian | 5 | 4.3 |
| Hispanic/Latino | 18 | 15.5 | Pescatarian | 2 | 1.7 |
| Other/Multiracial | 25 | 21.6 | Vegetarian | 0 | 0.0 |
| *Meat Dependence* |  |  |  |  |  |
| Mean (SD) | 4.03/5 (0.80) |  |  |  |  |

### 3.2. Consumer Sensory Attribute Ratings

Figure 2 and Table 3 show the mean sensory ratings on a 5-point Likert scale for Soy, Soy-Wheat, Beef, and Beef-Mushroom meatballs. The four meatballs showed significant differences in all eight sensory attributes (Friedman’s test; all *p* < 0.001). Effect sizes ranged from small (*W* = 0.12–0.24) to medium (*W* = 0.32–0.38), with hardness showing the strongest differentiation.

**Table 3:** Consumer sensory ratings and instrumental texture parameters. Sensory attributes rated on 1-5 scale; TPA parameters shown as mean ± SD. Values with different superscript letters within rows are significantly different (*p* < 0.05).

| Parameter | Soy | Soy-Wheat | Beef | Beef-Mushroom |
| --- | --- | --- | --- | --- |
| *Consumer Sensory Ratings (1-5 scale)* |  |  |  |  |
| Chewiness | 2.58<sup>a</sup> | 3.79<sup>b</sup> | 3.44<sup>b</sup> | 3.05<sup>ab</sup> |
| Hardness | 2.36<sup>a</sup> | 3.56<sup>b</sup> | 3.03<sup>b</sup> | 2.91<sup>ab</sup> |
| Moistness | 2.78<sup>ab</sup> | 2.04<sup>a</sup> | 3.25<sup>b</sup> | 3.27<sup>b</sup> |
| Fibrousness | 2.77<sup>a</sup> | 3.28<sup>ab</sup> | 3.43<sup>b</sup> | 3.08<sup>ab</sup> |
| Meatiness | 2.77<sup>a</sup> | 2.72<sup>a</sup> | 4.41<sup>b</sup> | 3.87<sup>b</sup> |
| Fattiness | 2.43<sup>a</sup> | 2.07<sup>a</sup> | 3.40<sup>b</sup> | 3.42<sup>b</sup> |
| Tastiness | 2.75<sup>a</sup> | 2.34<sup>a</sup> | 3.46<sup>b</sup> | 3.51<sup>b</sup> |
| Softness | 3.21<sup>b</sup> | 2.42<sup>a</sup> | 3.03<sup>ab</sup> | 3.09<sup>b</sup> |
| *Instrumental TPA Parameters* |  |  |  |  |
| Hardness (N) | 1.14 ± 0.22<sup>b</sup> | 1.41 ± 0.60<sup>b</sup> | 2.31 ± 0.78<sup>a</sup> | 2.56 ± 0.69<sup>a</sup> |
| Stiffness (kPa) | 45.4 ± 8.8<sup>b</sup> | 56.2 ± 23.9<sup>b</sup> | 91.7 ± 30.9<sup>a</sup> | 101.7 ± 27.5<sup>a</sup> |
| Cohesiveness | 0.29 ± 0.05<sup>b</sup> | 0.32 ± 0.07<sup>b</sup> | 0.43 ± 0.09<sup>a</sup> | 0.43 ± 0.06<sup>a</sup> |
| Springiness | 0.27 ± 0.06<sup>b</sup> | 0.34 ± 0.09<sup>b</sup> | 0.55 ± 0.11<sup>a</sup> | 0.52 ± 0.05<sup>a</sup> |
| Resilience | 0.22 ± 0.03<sup>b</sup> | 0.27 ± 0.04<sup>b</sup> | 0.37 ± 0.04<sup>a</sup> | 0.40 ± 0.07<sup>a</sup> |
| Chewiness (N) | 0.09 ± 0.03<sup>b</sup> | 0.19 ± 0.17<sup>b</sup> | 0.56 ± 0.26<sup>a</sup> | 0.57 ± 0.19<sup>a</sup> |

**Figure 2:**
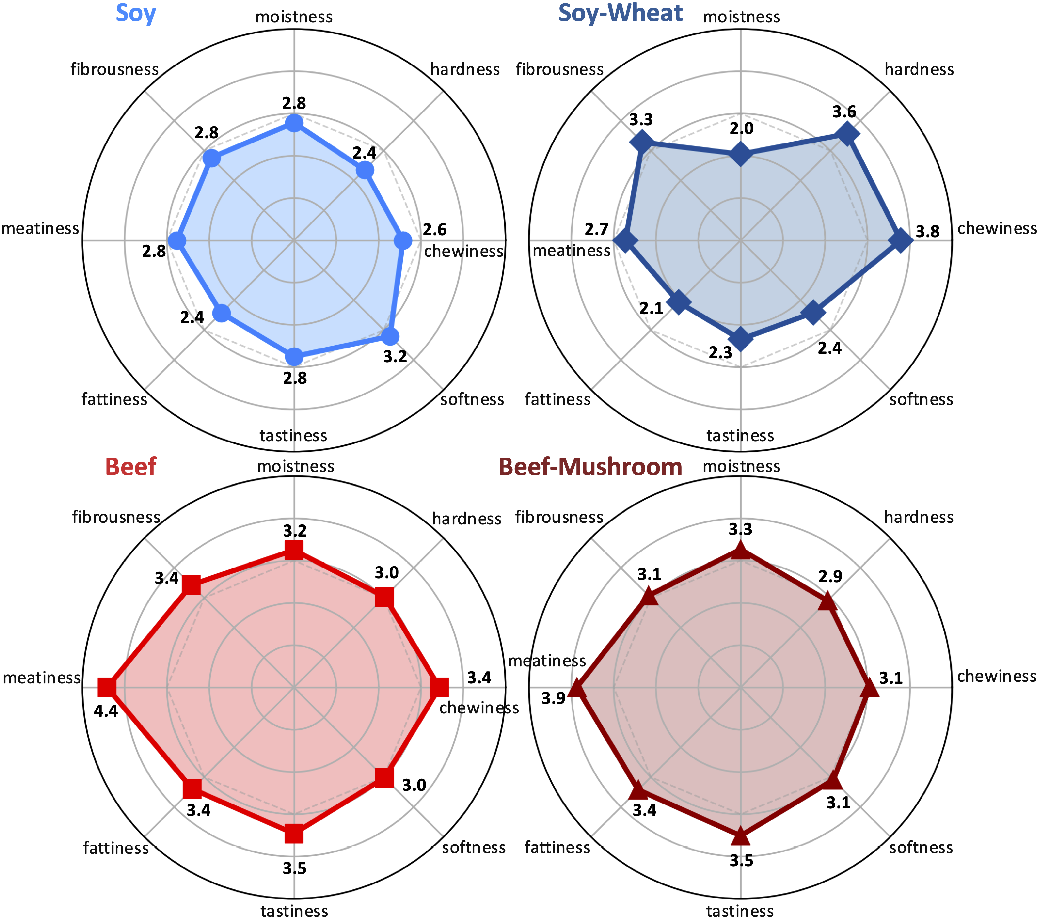
Individual sensory profiles of plant-based and animal-based meatballs. Radar charts display mean consumer ratings (1–5 scale) for eight sensory attributes—moistness, hardness, chewiness, softness, tastiness, fattiness, meatiness, and fibrousness—for Soy (top left), Soy-Wheat (top right), 100% Beef (bottom left), and Beef-Mushroom (bottom right) meatballs. Values represent mean ratings across all participants (*n* = 116).

The Beef meatball received the highest overall ratings across all sensory categories: moistness, hardness, chewiness, softness, tastiness, fattiness, meatiness, and fibrousness. Beef-Mushroom scored similarly in almost all categories, except for meatiness, where it scored lower (4.4 vs. 3.9), although not significantly so. In contrast, Soy and Soy-Wheat meatballs received significantly lower scores for meatiness, fattiness, and tastiness with a gap of approximately 1 full point. Soy-Wheat was the highest rated for chewiness and hardness, while Soy was the highest rated for softness. Post-hoc pairwise comparisons revealed substantial sensory differences between the two plant-based products: Soy-Wheat was rated significantly harder (3.6 vs. 2.4; *p* < 0.001) and chewier (3.8 vs. 2.6; *p* < 0.001) than Soy.

Figure 3 compares participants’ ratings for animal vs. plant meatballs across all eight sensory categories. We calculated each participant’s difference score as Δ = (*Soy* + *Soy-Wheat*) - (*Beef* + *Beef-Mushroom*). Chewiness shows a balanced distribution of ratings with 41% of participants rating the animal meatballs chewier and 41% rating the plant meatballs chewier; 18% of participants have no difference in scores. Hardness, softness, and fibrousness also have mixed scores between higher animal and higher plant and large percentages of participants rating no difference at 25%, 26%, and 22%, respectively. In contrast, moistness, meatiness, fattiness, and tastiness all skew strongly toward higher animal scores than plant, with 71%, 83%, 84%, and 77% of participants giving the animal meatballs higher scores, respectively. These sensory categories also have the lowest proportion of participants rating no difference between the animal and plant meatballs at 11% or under.

**Figure 3:**
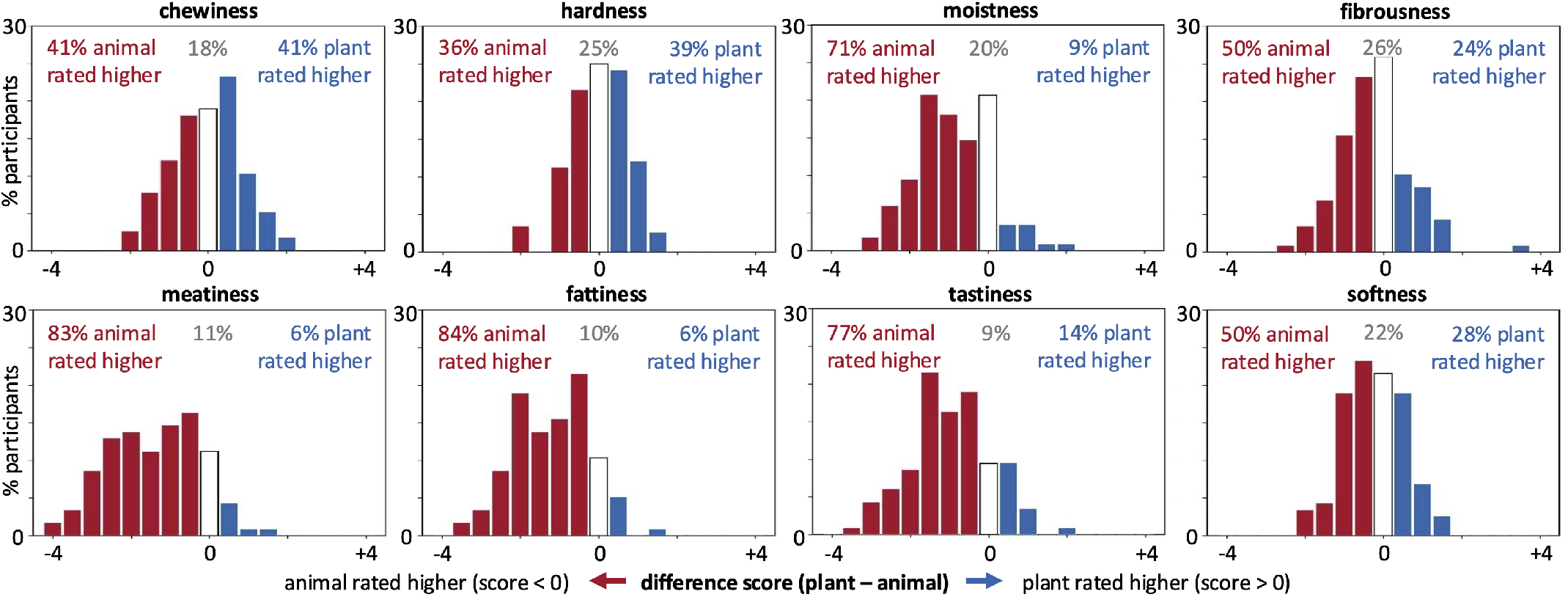
Distribution of differences in sensory rankings for animal vs. plant meatballs. The difference score for each participant (*n* = 116) is the sum of their Soy and Soy-Wheat scores minus the sum of their Beef and Beef-Mushroom scores for each sensory attribute. Red bars indicate higher scores for animal meatballs, blue bars indicate higher scores for plant meatballs, and the white bar indicates no difference in scores between animal and plant meatballs. Annotations show the total percentage of people in each category: higher animal scores (red), no difference (grey), higher plant scores (blue).

### 3.3. Just-About-Right Penalty Analysis

Figure 4 shows the Just-About-Right sensory profiles for Soy, Soy-Wheat, Beef, and Beef-Mushroom meatballs. The diverging bar charts reveals when an attribute, fibrousness, fattiness, savoriness, chewiness, or moistness is either “not enough” or “too much.” The percentage of respondents who rated each attribute “just about right” is listed above each bar. A perfect meatball would have a “just about right” rating of 100% represented by a line down each chart.

**Figure 4:**
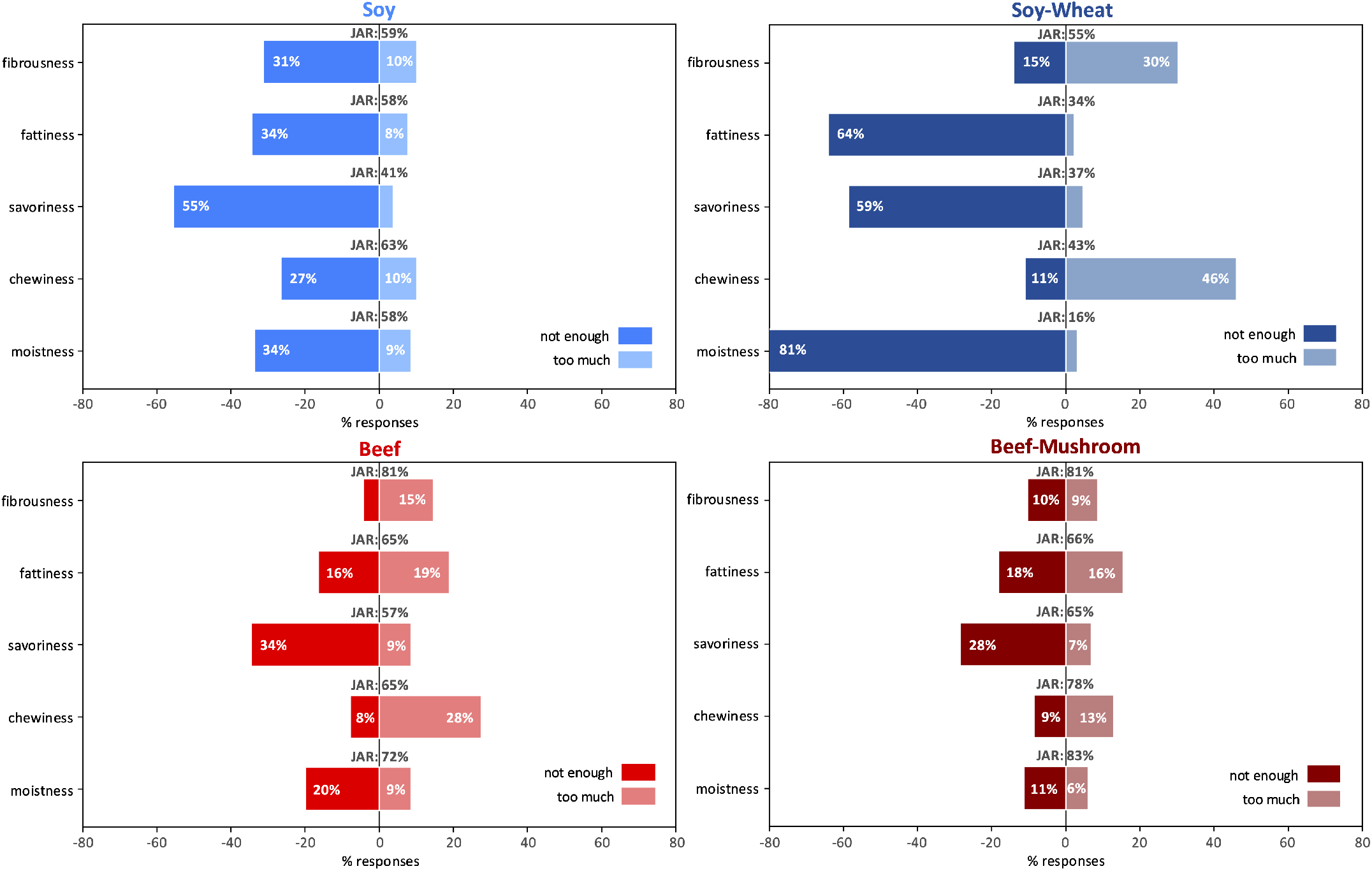
Just-About-Right (JAR) sensory attribute profiles for plant-based and animal-based meatballs. Diverging bar charts display the percentage of consumer responses (*n* = 116) rating each sensory attribute as “not enough” (left) or “too much” (right) for Soy (top left), Soy-Wheat (bottom left), Beef (top right), and Beef-Mushroom (bottom right) meatballs. The JAR percentage (responses rating the attribute as “just about right”) is annotated at the center of each bar.

The Soy-Wheat meatball has the largest absolute deviations from “just about right” across fattiness, savoriness, chewiness, and moistness, with fattiness, savoriness, and moistness being rated “not enough” and chewiness “too much.” Soy has the largest absolute deviation in fibrousness with 31% of responders rating it “not enough.” Soy is rated across-the-board as having predominantly not enough of all sensory attributes, while the three other meatballs have a mix between not enough and too much for different attributes.

Insufficient savoriness was penalized across both plant- and animal-based meatballs, with tastiness penalties ranging from 0.71 for Soy-Wheat to 1.36 for Beef, indicating that respondents who rated savoriness as “not enough” gave tastiness scores approximately 0.7–1.4 points lower than those who found savoriness “just about right.” Soy-Wheat showed bidirectional chewiness penalties for both “not enough” and “too much”, suggesting inconsistent texture perception across consumers. The Beef meatball exhibited the largest penalties overall, indicating that while it received higher absolute tastiness ratings, deviations from optimal attribute levels had a proportionally greater negative impact on its acceptance.

### 3.4. Texture profile analysis

To assess the quantitative mechanical attributes of the four meatballs, we tested 8 mm diameter by approximately 10 mm high samples with double compressions up to 50% strain at 25%/s (Figure 5, left). All meatballs showed highly reproducible force-time curves with relatively small standard deviations between samples. The Beef-Mushroom and Beef meatballs exhibit qualitatively similar curves, in shades of red, as do the Soy-Wheat and Soy, in shades of blue.

**Figure 5:**
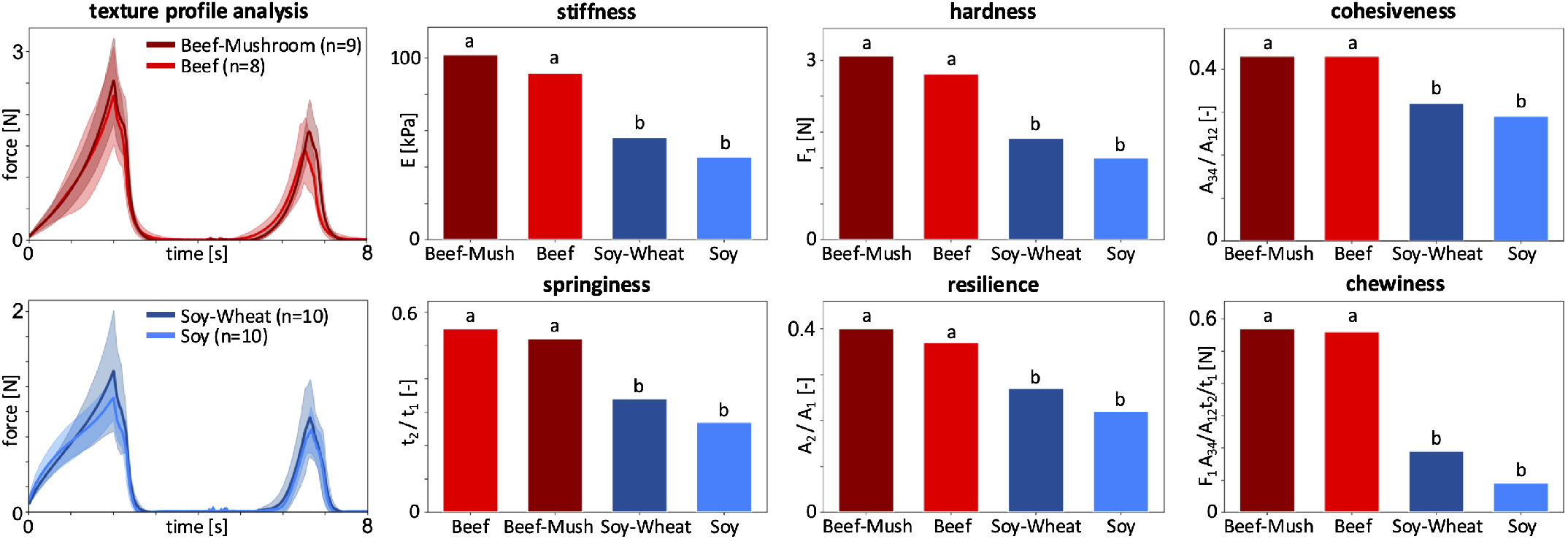
Texture profile analysis force-time curves and resulting parameters. Left plot: Force-time curves from double-compression tests at a loading rate of 25%/s up to 50% strain Beef, Beef-Mushroom, Soy-Wheat, and Soy meatballs (*n* = 37). Curves and shaded regions represent the mean and standard deviations for each meatball type. Bar plots represent mean stiffness, hardness, cohesiveness, springiness, resilience, and chewiness from force-time curves. Different letters annotated above the bar plots denote significantly different means with *p* < 0.05.

Using texture profile analysis, we extract stiffness, hardness, cohesiveness, springiness, resilience, and chewiness from the force-time curves. The bar plots confirm the visual similarity between the Beef-Mushroom and Beef and between the Soy-Wheat and Soy (Figure 5, right). Beef-Mushroom is the stiffest meatball at 101.7 kPa, then Beef at 91.7 kPa, followed by Soy-Wheat at 56.2 kPa, and Soy at 45.4 kPa (Tab. 3). Chewiness, which is the product of hardness, cohesiveness, and springiness, reveals the largest gap between meatballs: Beef-Mushroom has a chewiness of 0.57 N, Beef 0.56 N, Soy-Wheat 0.19 N, and Soy 0.09 N. Notably, the two beef products in dark red and light red are nearly two times higher in stiffness, hardness, and springiness, and almost three times higher in chewiness, than the two plant-based products in dark blue and light blue. In fact, statistical analysis reveals that for all six texture parameters, the two beef meatballs are significantly different than the two plant meatballs (*p* < 0.05), but there are no differences within each category of beef vs. plant.

### 3.5. Texture-Sensory Correlation Analysis

To examine the relationship between consumer sensory perceptions and instrumental texture measurements, we computed Kendall’s rank correlations, *τ*, between the eight sensory attribute ratings (Figure 2) and the six texture profile analysis parameters (Figure 5). A value of *τ* = 0 indicates that instrumental measurement product rankings do not correlate with sensory ratings, a value of *τ* = − 1 indicates perfect negative correlation, and a value of *τ* = +1 indicates perfect positive correlation. Direct comparisons between sensory and instrumental measures of the same attribute revealed no monotonic relationships (Figure 6): Mechanical hardness did not predict sensory hardness (*τ* = 0.00, *p* = 1.00), and mechanical chewiness did not predict sensory chewiness (*τ* = 0.00, *p* = 1.00). This disconnect was most apparent when comparing across categories: the two beef meatballs had approximately twice the instrumental hardness of plant-based samples (2.3-2.6 N vs 1.1-1.4 N), yet consumers did not perceive a proportional difference in sensory hardness ratings.

**Figure 6:**
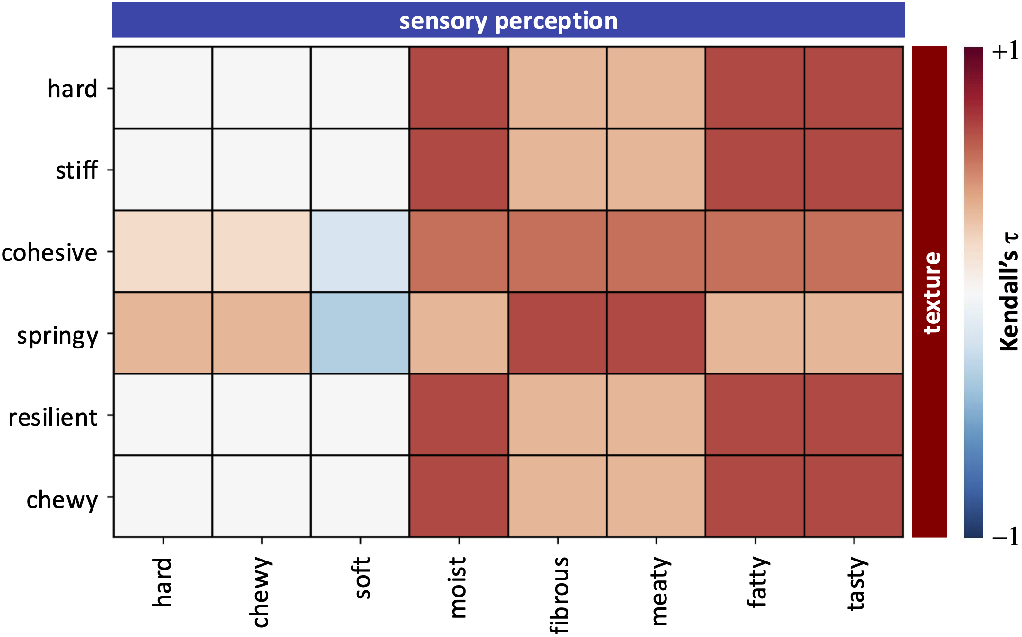
Kendall’s *τ* rank correlations between texture profile analysis and sensory perception parameters. Kendall’s rank correlation, *τ*, is computed for the ordered results from Figures 2 and 5 of Beef, Beef-Mushroom, Soy, and Soy-Wheat meatballs ranging from dark red, *τ* = +1, perfect positive correlation, to dark blue, *τ* = −1, perfect negative correlation.

In contrast, most texture parameters showed moderate-to-strong positive associations with perceived meatiness (*τ* = +0.33 to +0.67). Hardness, stiffness, springiness, and resilience each showed *τ* = +0.67 with meatiness, indicating that in 4 of 6 pairwise product comparisons, higher instrumental values corresponded to higher meatiness ratings. Texture parameters showed similarly consistent associations with perceived fattiness and tastiness (*τ* = +0.55 to +0.67). However, none of these correlations reached statistical significance. With only *n* = 4 products, Kendall’s *τ* requires perfect monotonic agreement (|*τ*| = 1.0) to achieve p < 0.05. The strong Pearson correlations reported in exploratory analyses (r > 0.90) likely reflect the influence of the large gap between animal and plant-based products rather than a robust monotonic trend across all four products. These findings should be interpreted as suggestive patterns and require replication with larger sample sizes to confirm.

### 3.6. Choice Factor Importance

Participants rated how eight different factors influence their choices between animal and plant-based meats in the dining hall on a scale between 0-100 (Figure 7). The participants prioritized these dining hall choice factors significantly differently (Friedman *χ*^2^(7) = 313.07, *p* < 0.001, Kendall’s *W* = 0.349. Flavor and texture are rated as the most important factors when choosing between animal and plant-based options with mean importance scores of (82/100) and (65/100). Familiarity/comfort, health/nutrition, and availability all received moderate average importance scores at (60/100), (53/100), and (50/100). Interestingly, climate sustainability (28/100), animal welfare (25/100), and social/cultural norms (24/100), are rated much lower than the other five factors.

**Figure 7:**
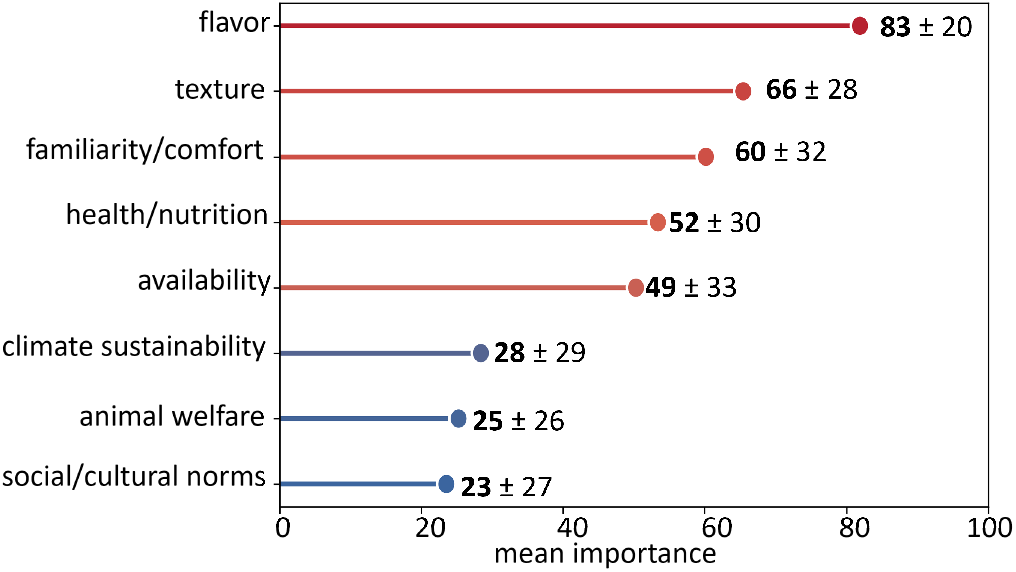
Factors influencing the choice between animal and plant-based meat in the dining hall. Participants (*n* = 116) rated the importance of eight factors (flavor, texture, familiarity/comfort, health/nutrition, availability, climate sustainability, animal welfare, social/cultural norms) using a sliding-scale [0-100] for each factor. The mean importance rating is plotted from highest to lowest importance factor.

### 3.7. Demographic and Individual Difference Analyses

Meat attachment scores and tastiness ratings were not significantly correlated for any product (all *p* > 0.05). Males rated Soy-Wheat significantly higher than females (*U* = 2496.0, *p* = 0.015, *r* = 0.24). We did not find any other gender differences.

Ethnicity did not significantly correlate with tastiness ratings of any product (Kruskal-Wallis, all *p* > 0.35). Diet type (omnivore vs. non-omnivore) did not significantly predict tastiness ratings (all *p* > 0.30). Neither plant-based dining frequency nor self-reported percentage of plant-based diet correlated with tastiness ratings (all *p* > 0.20). Choice factor importance ratings (flavor, texture, health, sustainability, animal welfare, familiarity, availability) showed minimal correlations with sensory attribute ratings (Spearman |*ρ*|< 0.25, 7.8% significant at p < 0.05), indicating that food choice priorities did not systematically bias their sensory evaluations.

### 3.8. Qualitative Findings

We identified common trends in participant comments using thematic analysis (*n* = 98 participants provided comments):

The themes mentioned most frequently were flavor/taste (*n* = 39 responses), specific dish suggestions (*n* = 37), texture concerns (*n* = 15), tofu preference (*n* = 14) and meat-like quality (*n* = 12). The sentiment was predominantly constructive, with participants offering suggestions for product improvement and dining hall variety.

## 4. Discussion

This study provides a comprehensive comparison of two plant-based and two animal-based meatballs using both consumer sensory evaluation in a naturalistic university dining setting and instrumental texture analysis in the lab. The findings reveal persistent sensory gaps between product categories while identifying specific improvement priorities for both plant-based and animal-based formulations.

### Animal meatballs outperform plant meatballs in key sensory attributes

The meatiness gap (Δ = 1.40 on a 5-point scale, *p* < 0.001) reveals the largest difference, consistent with evidence that meatiness replication remains the most challenging aspect of plant-based meat development (Kumari et al., 2024). Clear expectations set by “tastes like meat” marketing are often not fulfilled, particularly among flexitarian consumers (Fiorentini et al., 2023), a finding our data support given the predominantly omnivorous sample (89.8%). The moistness gap (Δ = 0.85) aligns with mechanistic research which shows that plant proteins lack the water-binding capacity of animal myofibrillar proteins (McClements & Grossmann, 2021). For example, in a recent blinded restaurant survey, a bean burger received significantly lower moisture ratings than the Big Mac (9% vs. 32%) and was perceived as significantly more dry (51% vs 18%) (Tac et al., 2026). Moisture release during mastication differs fundamentally between soy-based and meat-based products due to differences in protein network structure (Sha & Xiong, 2020). Our Just-About-Right data show that the two plant-based meatballs were perceived as less moist compared to the two animal-based meatballs (all *p* < 0.001).

### Texture has a variable impact on consumer acceptance

Substantial variation existed within the plant-based category. Soy and Soy-Wheat differed dramatically in texture perception, with Soy-Wheat rated significantly harder and chewier (effect size *r* = 0.76). This heterogeneity reflects the diverse formulation approaches in the market: Soy uses heme-containing soy leghemoglobin with high-moisture extrusion, while Soy-Wheat employs a soy-wheat protein blend with different processing parameters (He et al., 2020). These results caution against treating “plant-based meat” as a monolithic category in sensory research. The two animal products, despite differing in composition (Beef vs. Beef-Mushroom), did not differ significantly on most consumer measures. This finding has practical implications for Beef-Mushroom meat products that seek to reduce meat content while maintaining consumer acceptance, consistent with evidence that mushroom-beef blends up to 30% substitution maintain sensory equivalence to plain beef patties (Mylan et al., 2023).

### Mechanical features fail to predict sensory features

Consumer-perceived chewiness and hardness did not differ significantly between plant and animal categories despite substantial instrumental differences. This finding parallels recent work which found that trained panelists reported smaller differences in plant and animal texture features than instrumental measures (Bakhsh et al., 2021). This disconnect suggests that consumers integrate multiple sensory inputs beyond mechanical resistance—including moisture release, particle break-down, and temperature-dependent changes during mastication. Consumer ratings showed no category difference for perceived chewiness or hardness. Exploratory correlation analysis quantified this disconnect: TPA hardness did not predict sensory hardness (Kendall’s *τ* = 0.00, *p* = 1.00), and TPA chewiness did not predict sensory chewiness (*τ* = 0.00, *p* = 1.00). The Soy-Wheat product exemplified this paradox, ranking among the lowest in instrumental hardness (1.41 N) yet highest in consumer-perceived hardness (3.56/5). Strikingly, while TPA parameters failed to predict their corresponding sensory attributes, they showed consistent positive associations with perceived meatiness. TPA springiness, cohesiveness, chewiness, and hardness all showed Kendall’s *τ* = +0.67 with meatiness ratings, indicating that in most pairwise product comparisons, higher instrumental values corresponded to higher meatiness perception. While further studies with other products are needed to increase correlation power, this pattern suggests that mechanical texture properties contribute to the holistic perception of “meat-likeness” rather than mapping directly onto discrete sensory descriptors.

### None of the meatballs were savory enough

JAR penalty analysis revealed that insufficient savoriness penalized tastiness ratings across all four products (penalties 0.71–1.36 points), including both animal-based formulations. The savoriness deficit was actually larger for the Beef meatball (penalty = 1.36) than for either plant-based product. For dining hall operations, this implies that sauce accompaniments, seasoning adjustments, or umami-enhancing ingredients (mushrooms, fermented soy, nutritional yeast) could benefit all meatball offerings. These changes may be more actionable than protein-source modifications.

### Flavor and texture win over sustainability

Participants prioritized flavor (mean importance = 83/100) and texture (66/100) over sustainability (28/100) and animal welfare (25/100). This hierarchy aligns with evidence that while ethical concerns increase willingness to try alternative proteins, sensory properties remain the primary determinant of repeat consumption (Onwezen et al., 2021). These findings have direct implications for university food service strategy. Students respond more favorably to dishes presented with taste-focused labels than to dishes presented with health-focused labels. The relatively low importance ratings for sustainability and animal welfare suggest that ethical messaging alone is unlikely to drive adoption among this population.

### Limitations

Several limitations warrant consideration. The study population is homogeneous (96% aged 18–24, predominantly Stanford undergraduate students) which may limit generalizability to broader food service populations. The real-world dining setting introduces uncontrolled variability (accompaniments, hunger state, social context, self-selection bias) that laboratory studies avoid, although this design choice prioritizes ecological validity. The labeling with green and red flags maintains product anonymity but does signal protein category, which may have introduced expectation bias. Texture profile analysis at room temperature, rather than serving temperature, could affect absolute values, although relative comparisons likely remain valid. Finally, the cross-sectional design cannot address whether repeated exposure would reduce the observed sensory gaps through familiarization effects.

### Implications

For plant-based product developers, our data suggest prioritizing sensory improvements in moistness and meatiness over other texture modifications. In contrast to our measured instrumental texture differences, our sensory results suggest that perceived texture parity has already been achieved in all four meatballs of our study. For institutional food service, three actionable recommendations emerge: i) enhancing savoriness across all meatball offerings through seasoning or sauce modifications, ii) positioning plant-based options by culinary attributes rather than meatless status, and iii) recognizing that sensory quality improvements are likely more effective than sustainability messaging for driving adoption among young adult diners.

## 5. Conclusion

This study suggests that plant-based meatballs remain distinguishable from animal-based counterparts in a university dining setting, with significant gaps in perceived moistness, meatiness, fattiness, and tastiness. Instrumental texture analysis confirmed differences in mechanical features, although these differences did not map one-to-one onto consumer-perceived features such as hardness or chewiness. Among young adult university diners, hedonic factors dominated food choice over ethical considerations. Insufficient savoriness emerged as a universal improvement target across all four products of this study. These findings provide specific guidance for both plant-based product reformulation and institutional food service operations that seek to implement healthier food options.

## 6. Data availability

De-identified sensory data, mechanical texture analysis results, and data processing scripts are publicly available at https://github.com/LivingMatterLab/AI4Food.

## Acknowledgments

We thank Andrew Mayne and the dining hall staff at Stanford University for their assistance with sample preparation and data collection. This work was supported by the Stanford Plant-Based Diet Initiative. In addition, SS was supported by the NSF Graduate Research Fellowship, by the Stanford DARE Fellowship, AK by the Stanford Doerr School of Sustainability Accelerator, NZ by the Bio-X Summer Undergraduate Research Program, and EK by a Stanford Bio-X Snack Grant, by the NSF CMMI grant 2320933, and by the ERC Advanced Grant 101141626.

